# Gut Microbiome Signatures of Yorkshire Terrier Enteropathy during Disease and Remission

**DOI:** 10.1101/2022.08.25.505284

**Authors:** Pavlos G. Doulidis, Alexandra I. Galler, Bela Hausmann, David Berry, Alexandro Rodríguez-Rojas, Iwan A. Burgener

## Abstract

The role of the gut microbiome in developing Inflammatory Bowel Disease (IBD) in humans and dogs has received attention in recent years. Evidence suggests that IBD is associated with alterations in gut microbial composition, but further research is needed in veterinary medicine. The impact of IBD treatment on the gut microbiome needs to be better understood, especially in a breed-specific form of IBD in Yorkshire Terriers known as Yorkshire Terrier Enteropathy (YTE). This study aimed to investigate the difference in gut microbiome composition between YTE dogs during disease and remission and healthy Yorkshire Terriers. Our results showed a significant increase in specific taxa such as *Clostridium sensu stricto* 1, *Escherichia-Shigella*, and *Streptococcus*, and a decrease in *Bacteroides, Prevotella, Alloprevotella*, and *Phascolarctobacterium* in YTE dogs compared to healthy controls. No significant difference was found between the microbiome of dogs in remission and those with active disease, suggesting that the gut microbiome is affected beyond clinical recovery.

## Introduction

Chronic or relapsing gastrointestinal (GI) signs are a hallmark of canine and human inflammatory bowel disease (IBD)^1,2^. Canine IBD is diagnosed after histologic evidence of a characteristic intestinal inflammation pattern and after excluding other systemic, endocrine, neoplastic and infectious causes of chronic GI signs^3,4^. The underlying causes of IBD are not fully understood, but it is thought to result from a combination of genetic factors, environmental exposures, intestinal immune reactivity, and gut microbiome composition^5^. The synergic effect of the genetic background, environmental factors, intestinal immune reactivity, and the intestinal microbiome creates pathomechanisms difficult to disentangle^6,7^. Despite ongoing research, our understanding of the mechanisms driving IBD remains limited. Studies in animal models have primarily relied on inducing GI inflammation through various experimental methods in mice^8^.

IBD in humans manifests in various clinical forms and can partly result from unbalanced crosstalk between gut luminal content and the mucosal immune system. IBD encompasses a continuum of human clinical disorders, ranging from Crohn’s disease through indeterminate colitis to ulcerative colitis. Some patients may exhibit significant clinical and histological overlap between these forms or even develop one form from another^9^. The development of human IBD is also influenced by a combination of genetic, environmental, and pathogenic factors^9^.

The gut microbiome comprises bacteria, archaea, viruses, and eukaryotic organisms that reside in the GI tract ^10,11^. The bacterial component is the largest and provides essential digestive functions, such as fermentation of fibres and production of short-chain fatty acids (SCFA)^12^ and indoles^13^. Firmicutes, Bacteroidetes and Fusobacterium are the phyla that co-dominate healthy canine gut microbiome^14^. Some bacterial genera, such as *Escherichia, Collinsella, Megamonas, Lactobacillus* and *Streptococcus*, are consistently found in faecal samples of healthy dogs, indicating the presence of a core faecal bacterial community^15^. Characteristic examples include Clostridia and Bacilli ^16^ of the phylum Firmicutes, some of which contribute to the production of SCFAs. Furthermore, members of the phylum Bacteroidetes, like *Prevotella* and *Bacteroides* show high but also variable abundances in the healthy canine gut microbiota^1^.

In general, human IBD entails changes in intestinal microbiome (dysbiosis), focal translocation/infiltration of bacteria across the mucosal barrier into deeper intestinal layers. There is also an altered mucosal response to bacterial invasion, development of chronic granulomatous inflammation and activation of adaptive immunity as a consequence of compensatory mechanisms to minor defects of innate immunity or autophagy^10^. Genetic factors can involve variants of different loci that provoke a leaky epithelial barrier and/or impair phagocytosis and autophagy. Independent of the particular combination of factors in each patient, the typical result is a vicious circle of dysbiosis and granulomatous inflammation mediated by the immune system^10^.

Canine IBD exhibits several pathophysiologic and phenotypic similarities with humans IBD, as well as some uniquetraits. While IBD can affect any dog breed, certain breeds may display breed-specific disease phenotypes. Inbreeding in modern dog is similar to geographically highly isolated human populations, providing an excellent opportunity to better explore disease pathomechanisms of various complex disorders, including IBD^17^. In Yorkshire Terriers, retrospective studies have described an enteropathy distinct from other breeds^18,19^, which suggests the existence of a breed-specific “Yorkshire Terrier enteropathy” (YTE). Besides the classic chronic GI signs, clinical symptoms such as low oncotic pressure and effusions along with laboratory findings as hypoalbuminemia, hypocalcaemia and hypomagnesemia have been shown to be more prevalent. Histologically, severe intestinal lymphatic dilation, crypt lesions, and villous stunting are features commonly described in patients with YTE^19^.

In a previous study, we reported alterations in the blood metabolic profile of a cohort of Yorkshire terrier dogs suffering from YTE. Our findings showed that plasma metabolite levels remained altered during remission and differed considerably from healthy controls^17^. Additionally, changes in the faecal microbiome and the faecal metabolic profiles lead to imbalances in bile acid metabolism, sterols, and fatty acids in the same patient group^11^. To better understand the pathomechanisms of YTE and to contribute to potential therapeutic advancements, this study aimed to investigate dysbiosis related to the disease and compare it with healthy controls and dogs that had entered remission in the same cohort. To do this, we have characterised the bacterial microbiome in fresh faecal samples to determine its composition and relationship with the animals’ clinical state. Our study is the first work that addresses the microbiome alteration of the breed-specific Yorkshire Terrier enteropathy in the context of IBD with the advantage of decreasing genetic variability observed in other multi-breed IBD research.

## Results

A total of 39 Yorkshire terrier dogs were enrolled in this study and divided into the two initial groups: 13 subjects with confirmed IBD composed the YTE group, while 26 healthy dogs were used as a control group (see eligibility and group separation in Fig. 1). The disease group (YTE) was comprised of 9 females, four of which were spayed, and 4 males, with two of them neutered. The mean age was 6.4 years ± 1.7 years (mean+sd), and the mean body weight was 3.7 kg ± 1.5 kg (mean+sd). Before enrolment, dogs were fed diverse commercial diets. The primary symptoms on presentation were a history of chronic or intermittent GI signs (N=10) followed by pleural and/or abdominal effusion (N=3). Mean faecal scores were 4.5 out of 7 ± 1.1, and body conditioning scores were 4.3 out of 9 ± 1. Muscle conditioning scores were A (from A to D) in nine dogs, B in one, C in two, and D in one dog. The severity of the YTE was classified as mild to severe according to canine chronic enteropathy activity index (CCECAI, mean 9.2 ± 3, Fig. 2). World Small Animal Veterinary Association (WSAVA) scores indicated mild or moderate histological changes in the duodenum (median 3.5, range 2–11), with a predominant lymphoplasmacytic infiltration of the intestinal mucosa in all dogs (N=13). Crypt lesions (N=5), villus blunting (N=4), lymphangiectasia (N=3) and additional eosinophilic (N=4) or neutrophilic (N=3) infiltration were also found.

**Fig. 1.**
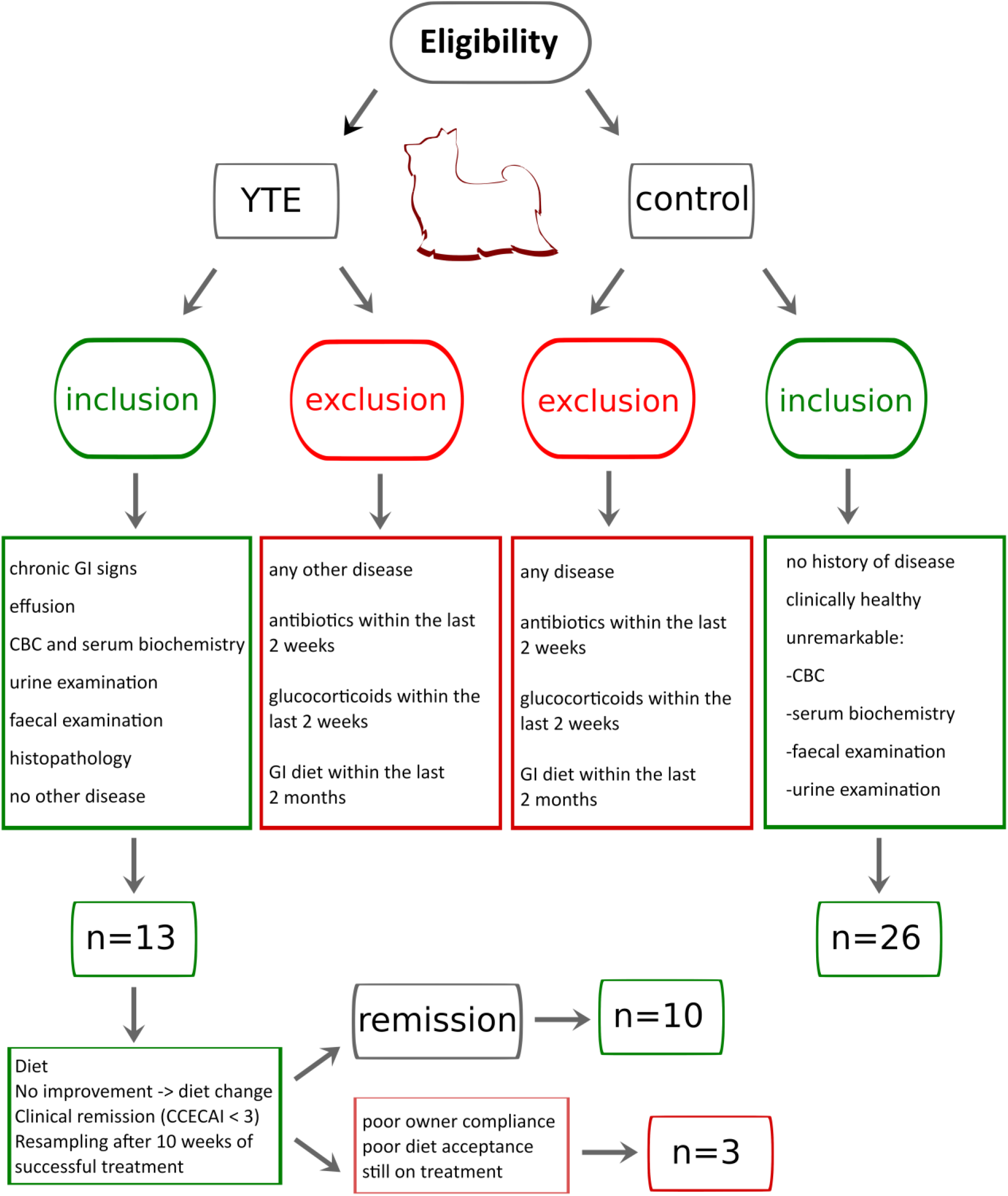
Inclusion and exclusion criteria for subject selection in this study. Diagram illustrating the inclusion and exclusion criteria applied to determine the composition of the three groups (YTE, control and remission).

**Fig. 2.**
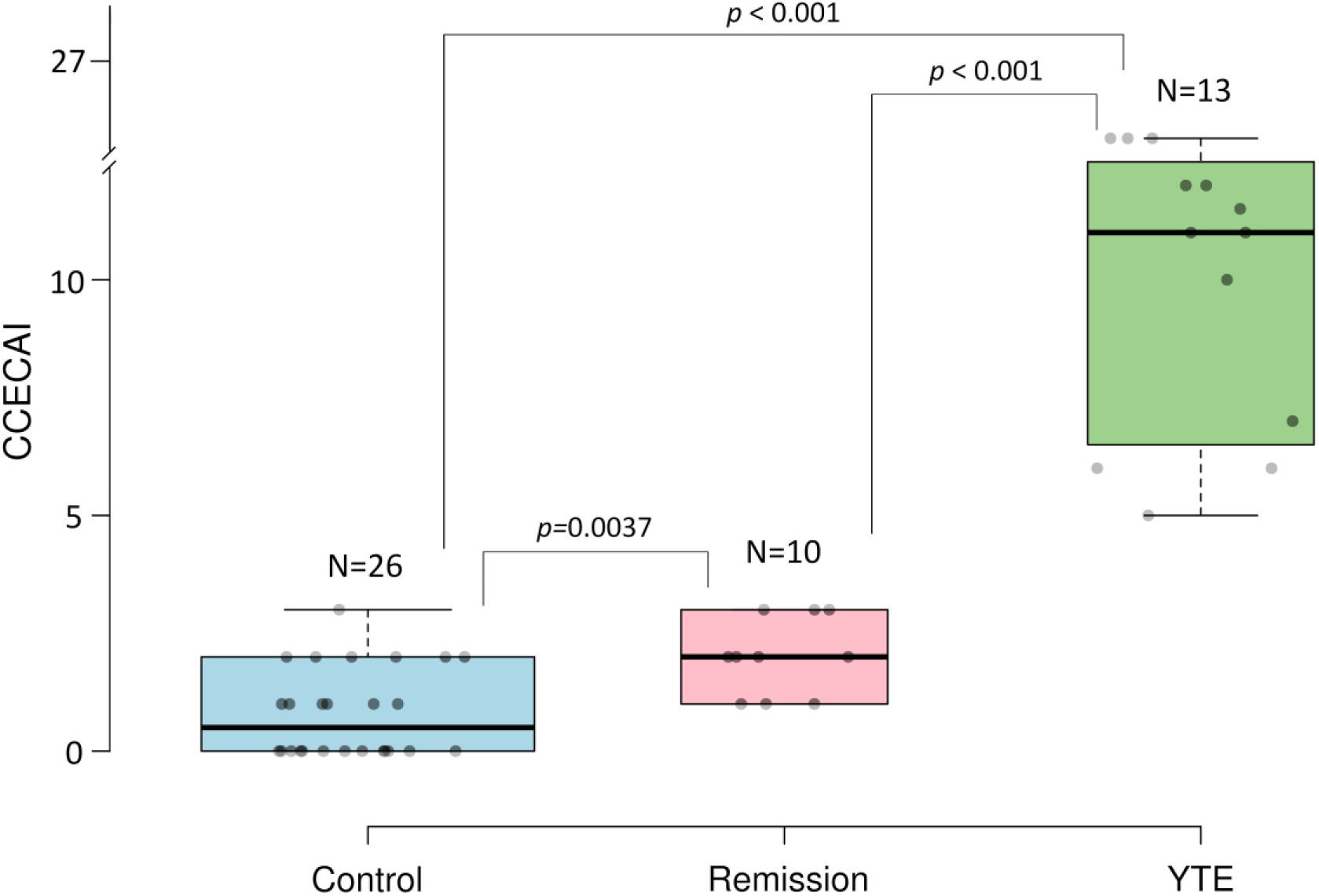
Classification of the Yorkshire terrier cohort used in this study according to the canine chronic enteropathy activity index (CCECAI). This index was used to calculate clinical severity of the enteropathy in all dogs. Assessing the severity of alterations in 9 different categories, including attitude and activity, appetite, vomiting, faecal consistency, defecation frequency, weight loss, serum albumin concentration, peripheral edema and ascites, and pruritus. Differences among groups were inferred from a Kruskal-Wallis test (*p*< .00001) followed by Mann-Whitney U-test to compare the three groups.

Ten dogs in the YTE group achieved clinical remission (CCECAI <3) and were resampled for gut microbiome analysis (Fig. 2). One dog was still under treatment at the time of resampling. Poor owner compliance led to the exclusion of one dog, and another dog was also excluded due to poor clinical response to the prescribed diets. From the 10 revaluated dogs, 8 were clinically well-controlled, received a hydrolysed diet, and were resampled 69–86 days (median 74 days) post-diagnosis. Two dogs reached clinical remission only after prednisolone administration and were resampled 126 and 134 days post-diagnosis (respectively 71 and 74 days after initiation of prednisolone). An improvement was noticed in faecal scores (mean 2 ± 0.5), body conditioning scores (mean: 4.7 ± 0.7) and muscle conditioning scores (A, N=8; B, N=1; C, N=1).

The control group (N=26) consisted of 14 females, of which 8 were spayed, and 12 males, of which 6 were neutered. All dogs were healthy Yorkshire Terriers. The mean age of the dogs was 8.2 ± 3.2 years. The mean body weight was 3.9 ± 1.35 kg. The mean faecal score was 2 ± 0.8, while the mean body conditioning score was 5.5 ± 0.8, and the muscle conditioning score was A (N=20) or B (N=6). Twenty-five dogs in the control group received diverse commercial diets (19 canned and 6 dry food). One dog was fed with a Bones and Raw Food (BARF) diet. Sex and body weight did not differ significantly between healthy and YTE dogs. Dogs in the healthy control group were older than dogs suffering from YTE (*p*=0.04). The median CECCAI score in the healthy control group was 0.5 (range 0-3), which was significantly lower compared to dogs with YTE (*p* < 0.001, Fig. 2).

As expected, statistical analysis of clinicopathological parameters showed poor predictive value (*p* > 0.05). A list of the analysed clinicopathological parameters is provided in the supplementary material (Supplementary Table S1). In addition, a principal component analysis that included all these clinical markers failed to segregate the three studied groups (YTE, Remission and Control, Supplementary Fig. S1).

Next, we determined the relative abundances of all microbial taxa from faecal material based on the number of 16 rRNA gene amplicon reads. Raw sequencing data is available in the NCBI BioProject database with the accession number PRJNA861113. Around 420 different bacterial taxonomic categories were detected. After filtering, we identified 468 unique ASVs and 159 unique genera. A complete table of identified microbiome bacterial genera is available in supplementary materials (Supplementary Table S2). The most dominant bacteria phyla sequenced were *Proteobacteria, Firmicutes, Fusobacteria and Bacteroides*.

The gut microbiome composition of the dogs in the YTE group did not show any difference in the alpha diversity of amplicon sequencing variants (ASVs, representing unique amplicon sequence types) (Fig. 3A), indicating that the alteration of the structure of the bacterial community concerning its richness was not statistically significant for the studied cohort. Nevertheless, the quantitative difference in the relative abundance of several taxa presented significant differences in beta diversity, indicating an imbalance in the abundance of particular microbes between healthy dogs and those with YTE or in remission (Fig. 3B and 3C). There were, however, no significant differences between YTE and Remission groups in this index, although there are significant differences in several species’ abundance that we detail later in this manuscript. These results were the first indication that the dogs in remission did not reach the expected eubiosis, although they achieved clinical remission over the course of more than two months of treatment.

**Fig. 3.**
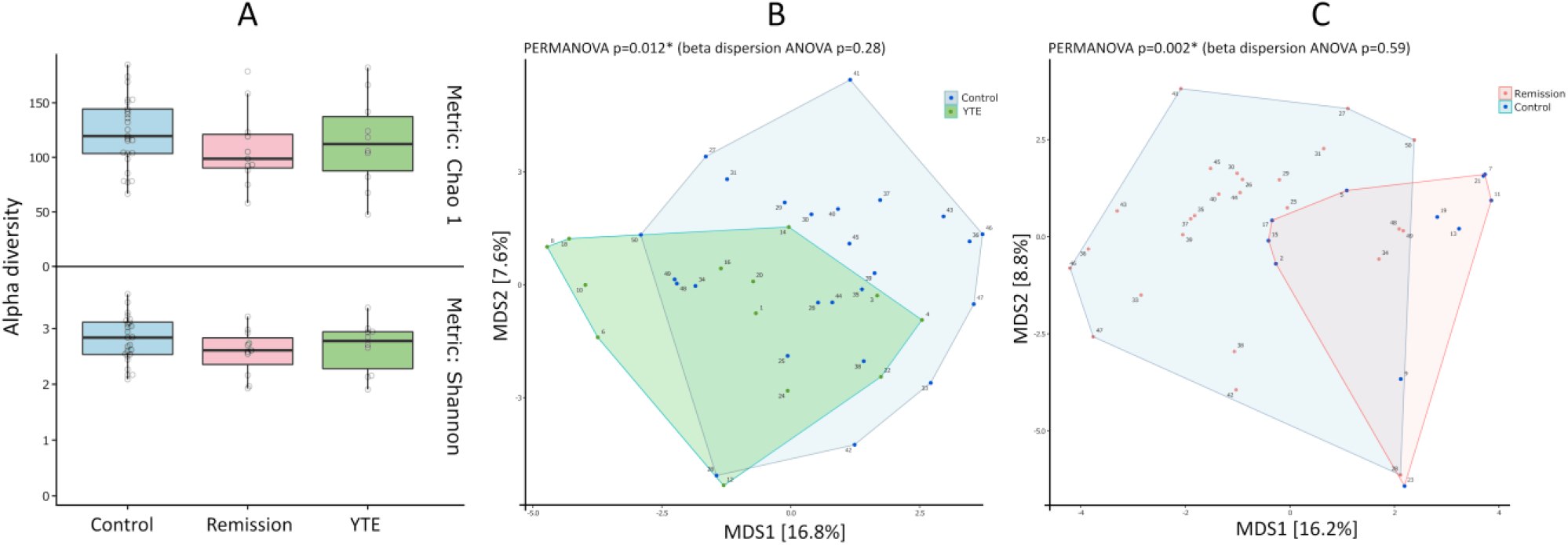
Alpha and Beta diversity analysis of microbiome of the studied Yorkshire terrier cohort. Alpha diversity of the microbiome in Yorkshire Terrier Enteropathy group (YTE) compared to control and remission groups (A): Shannon or Chao1 metrics show no significant differences among the three groups. Beta diversity shows significant difference between YTE and control (B) and between remission and control (C) but not between YTE and remission dogs.

We analysed the differentially abundant fraction expressed as numbers of reads of 16S rRNA gene. The taxonomic breakdown of the dominant ASVs across all samples can be seen in Supplementary Table S2 and Fig. S2. The quantitative analysis of the generic composition of the microbiome showed a significant decrease in the number of ASVs from the order Bacteroidales, mainly the genus *Bacteroides, Prevotella* and *Alloprevotella* (ranging between log2 fold-changes of 3 to 7) in YTE and remission dogs in comparison to the control. Selected bacteria that showed statistically significant altered abundance taking as reference the healthy dogs (control) are illustrated in Fig. 4. Other ASVs with decreased abundance belong to the genera *Catenibacterium, Fusobacterium, Succinivibrio* and *Helicobacter* (also with log2 fold-changes between 3 and 7) for YTE groups in comparison to healthy dogs. A more quantitative detailed differential abundance of bacteria with a significant fold-change is presented in Fig. 5. It is crucial to notice that remission dogs have more decreased bacterial ASVs than the YTE group in contrast to the control. If we compare the YTE group with remission, we observed the latter had a reduced number of ASVs belonging to *Peptoclostridium, Faecalibacterium, Ruminococcus, Cellulosilyticum, Fusobacterium, Clostridium, Streptococcus* (with log2 fold-change raging between 3 and 6). The other ASVs with lower abundance in remission compared to the control were classified to the genera *Peptoclostridium, Blautia*, and *Negativibacillus*, with a more moderate log2 fold-change of 2.

**Fig. 4.**
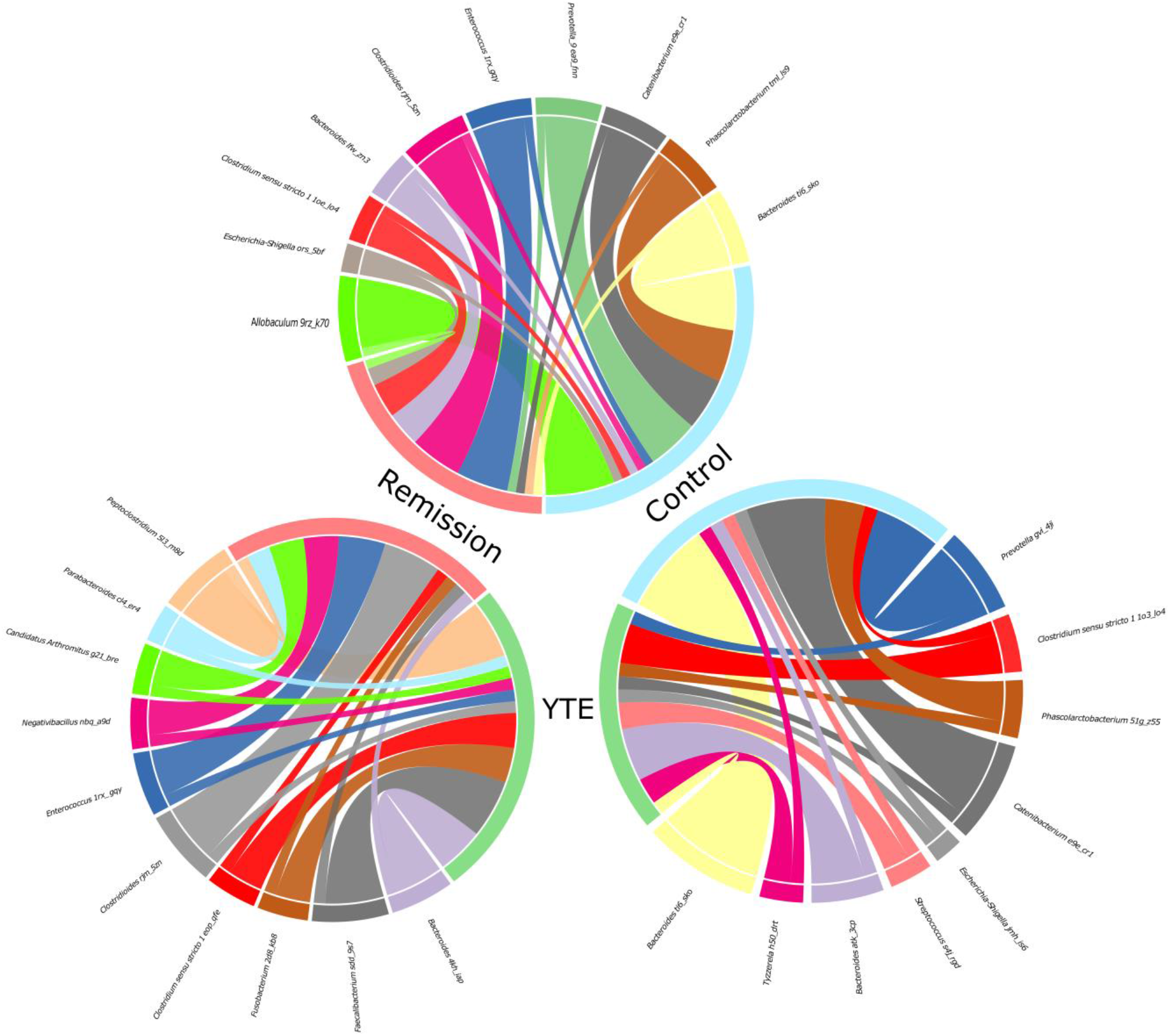
Chord diagrams showing the associations between differentially abundant taxa of Yorkshire Terrier Enteropathy (YTE), Remission and Control groups. Only the most abundant microbiome ASVs detected are shown. For better comprehension, each circle include two comparisons each time (remission in light red, YTE in green, and control in light blue). The chord diagrams show the key bacterial taxa identified by their comparative abundance. The outer ribbon identifies the respective clinical states (groups) and encompasses the perturbed taxa associated with each state. Chords connect taxa related to more than one state in the inner circle. Only significant hits are represented in these chord charts (at least *p*<0.05).

**Fig. 5.**
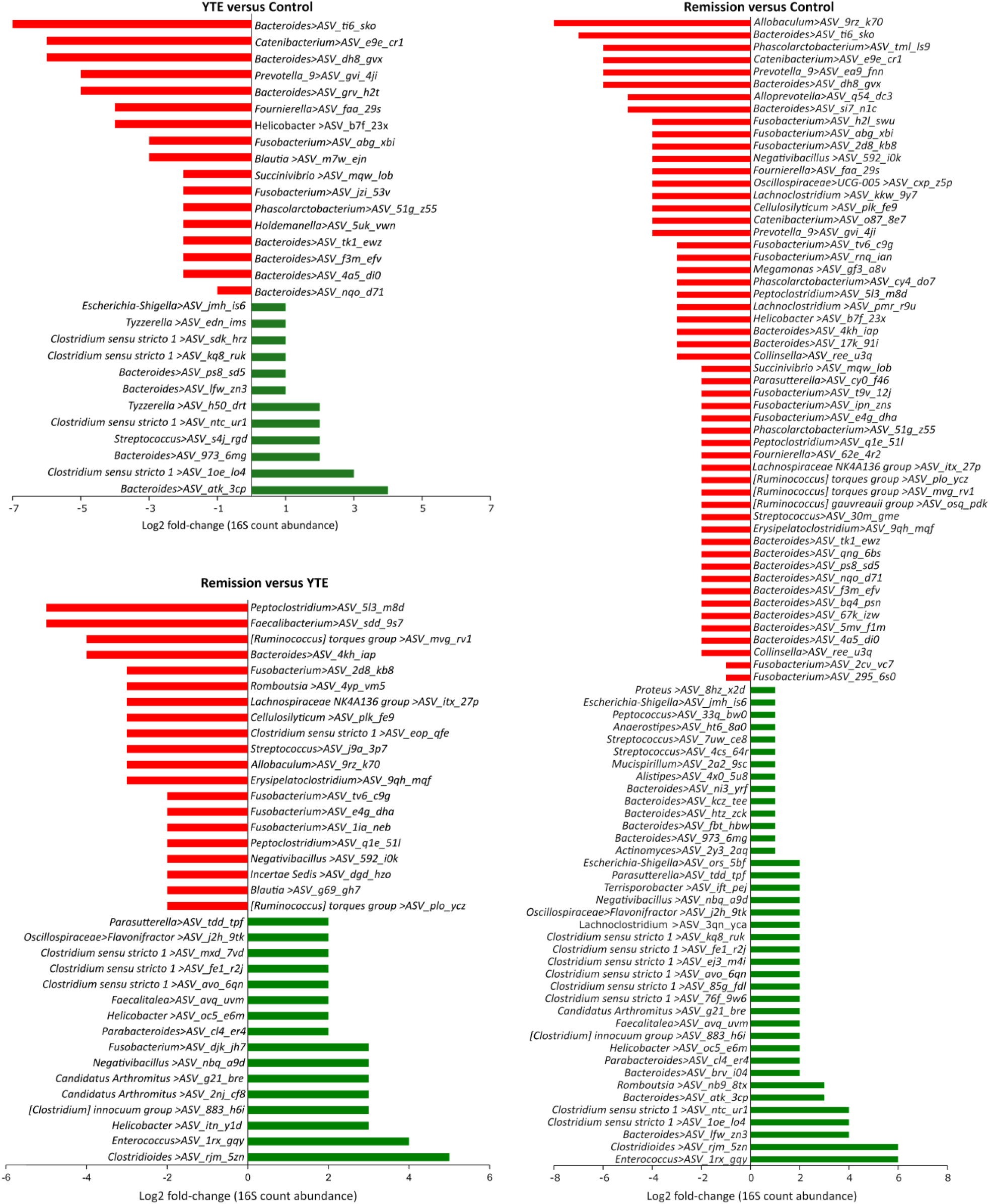
Differential abundances of the ASVs found in the YTE, Remission and Control group gut microbiomes. The shown data were determined by 16S rRNA gene sequencing. The red bars represent bacterial taxa showing a significant decrease during dysbiosis. In contrast, green bars represent those bacteria significantly more abundant in a given condition, expressed as the log2 fold-change in abundance. Only significant hits are represented in these plots (at least *p*<0.05).

At the same time, we found several ASVs with increased abundance in YTE and remission dogs compared to the control. Among the most increased ASVs for both groups, belong to the genera *Clostridium, Streptococcus, Tyzzerella, Candidatus Arthromitus*, and some *Bacteroides* from different species than those found in the groups of decreased bacteria (Fig. 5). All these ASVs were altered in abundance between log2 fold-change of 2 to 6. The same pattern observed in reduced bacterial populations is repeated here, where there is a higher number of enriched ASVs in the remission group compared to YTE. We observed a larger number of enriched ASVs, particularly belonging to the genera *Clostridium, Enterococcus, Romboutsia, Faecalitalea* and *Escherichia-Shigella*, among others, in remission dogs with a log2 fold-change between 2 to 6. The main difference in abundance between YTE and the remission group includes ASVs of the genera mentioned above that present higher quantities in the remission group of some bacteria, such as *Clostridiales, Enterococcus, Helicobacter, Negativibacillus, Fusobacterium* and *Candidatus* Arthromitus with a log2 fold-change between 3 to 6.

In summary, our dataset revealed a dysbiosis pattern that indicates a significant imbalance in the microbiome of several genera, both in terms of enrichment and depletion. Dogs in remission showed a greater disruption of the microbiome compared to the control group, with a larger number of hits of differentially abundant bacteria. The dysbiosis observed does not revert to balanced state, suggesting a long-term effect of the pathology, which clinical improvement as a result of the treatment, but suggesting also a different and new equilibrium.

## Discussion

The gut microbiome shows a marked individuality, and our understanding of the factors that imbalance gut microbial communities leading to pathogenic processes, has improved thanks to technological improvement and enhanced analysis capabilities^20^. Microbiome studies make an essential contribution and create intervention opportunities that will allow the development of personalised medicine in veterinary and human medicine. In our study, we analysed the gut bacterial microbiomes of a cohort of Yorkshire terriers suffering from IBD whose main advantage is the high degree of inbreeding that allows for minimising genetic variability that can mask other essential factors. Moreover, we examined the microbiome composition of sick dogs compared to controls and later reanalysed those dogs that entered remission. Here, we provide evidence that significant changes in the microbiome composition occur during this breed-specific YTE. Our data support previous studies that IBD dysbiosis varies depending on the studied cohorts.

The failure of clinical markers to predict signs of disease with the integration of all parameters in a principal component analysis (see Supplementary Fig. S1 and Table S1) confirmed how challenging the diagnosis of canine IBD is-as shown before in humans^21,22^. In addition, there is currently no cure for IBD, and a significant percentage of patients do not respond to clinical intervention^23^.

The gut microbiome profile in our Yorkshire terrier cohort is comparable to those reported in previous studies of other breeds. The same bacterial taxa show similar relative abundances in the gut microbiome. Although the microbiome member’s detection depends on the analysis methodology and definition, we have identified many similar genera. The lack of significance between YTE and Remission groups in alpha and beta diversity indexes indicates that perturbations of microbiome in IBD does not recover quickly. These results correlate well with a previous study of the metabolome in the same cohort^17^, where the metabolite profile did not recover to the previous status before the onset of the disease. These results also suggest the need of longer studies to find out long-term consequences of dysbiosis beyond clinically improvement. Additionally, we found an imbalance in bile acid metabolism, sterols, and fatty acids in the same patient cohort^11^. Reduced bile acid levels in the gut are associated with bacterial overgrowth and inflammation. Bile acids appear to be a significant regulator of the gut microbiota^24,25^. This relationship between the bile acid and gut bacteria suggests a mechanistic path on how microbiome alteration induces a malfunction of the GI tract that should be addressed in future studies. Bile acids are involved in the absorption of fat and the regulation of lipid homeostasis through emulsification. The gut microbiota is actively involved in the production of bile acid metabolites, such as deoxycholic acid, lithocholic acid, choline, and SCFAs such as acetate, butyrate, and propionate. Metabolites derived from the gut microbiota or modified gut microbiota metabolites contribute significantly to host pathophysiology. These alterations are thought to be a related with some GI cancers^26^.

The decreased abundance of several members of the *Bacteroidetes*, such as ASVs belonging to the genera *Bacteroides* and *Prevotella*, detected in this study have been previously reported in IBD in dogs and human^27,28^. Increase in other bacterial types such as *Clostridium, Escherichia-Shigella* and *Streptococcus* has been also previously reported^29^. In our study, we found that many bacteria of the *Fusobacterium* taxa are significantly decreased in YTE and Remission dogs. In contrast to humans, *Fusobacterium* is one of the genera typically associated with health in canines and has been proven to be very severely decreased in GI disease and after antibiotic treatment, also showing a slower recovery than other phyla^15,30^. Therefore, an increase of the *Fusobacteria* abundance through specific food ingredient supplementation may be a future therapeutic goal, as suggested before^15^. However, in our data, we found that dogs in remission have more hits for both decreased and increased ASVs in comparison to the control than the YTE groups. This could indicate a remodelling of the microbiome, in part due to the treatment, where other bacterial groups could occupy new available niches and suggests that perturbations in microbiota are difficult to return to previous state. In addition, this result also points to the fact that even though microbes are involved in the IBD pathogenesis, dogs showing clinical improvement have a wider perturbation of the microbiome. Thus, the microbiome may not be necessarily the trigger of IBD, but a consequence of the pathology as suggested elsewhere^31^.

The main limitation of our study is the small cohort of dogs with YTE and in remission. Another limitation is that gastroduodenoscopy was not repeated in dogs in clinical remission. Since gastroduodenoscopy in dogs has to be performed under general anaesthesia, it is not routinely used in the follow-up of canine IBD, and due to the relative invasiveness and risk, it would be ethically questionable to perform. This ethical consideration and owner consent are limitations that will not easily be overcome in client-owned dogs. In the future, canine intestinal organoids from affected dogs may allow us to study underlying pathobiological mechanisms and evaluate the impact of different therapeutic strategies *in vitro*^32^. Furthermore, dogs were not fed a standardised diet before presentation, which might have influenced microbial composition of the faeces at the beginning of the study. However, at the time of enrolment, all patients received comparable commercially available diets (except one dog with a BARF diet). Additionally, one dog in the treatment group received prednisolone because he did not respond well to the dietary treatment trial alone and later on entered in remission. Although one study reported no significant alterations in microbiome composition in a small cohort of dogs that received oral prednisolone for 14 days^33^, an effect cannot yet be completely excluded.

Despite the fact that the analysis of the microbiome of various species and environments using next-generation sequencing techniques has significantly improved our understanding of the metabolic, physiological, and ecological roles of microorganisms, the analysis is influenced by experimental conditions, complex downstream analysis, and sample collection among other factors^34^. We overcame these limitations, as much as possible, by adhering to our study to the best possible practices for sample collection and analysis in our conditions.

In this study, we could not prove a microbiome recovery to a healthy state, but we still think that gene changes and alterations in microbial interactions-networking might be present. Studies have proven the potential of network analysis uses in medicine^35,36^. Future studies on microbiome assembling rules might bring more light into the pathogenesis of canine IBD and the role that the gut microbiome plays in it. Different pathologies can influence the equilibrium of the microbiome, and in consistence with our study, restoration of the gut microbiome is incomplete after months of sickness. It is known that different gastrointestinal diseases can induce long-term changes in dog’s microbiome^37^. These durable perturbations of gut microbial composition are not exclusive to dogs. For example, a recent study showed that the human gut microbiome is not fully recovered three months after recovery from Coronavirus disease 2019 (COVID-19), which is known that causes alteration of gut microbiome alterations^38^. Antibiotics are also a known cause of dysbiosis. Another study indicates a prolonged or permanent switch to a stable alternative microbiome in the human gut microbiome, whose consequences remain unknown^39^.

## Conclusions

In summary, our analysis has shown that dogs with YTE show gut dysbiosis that did not alter the alpha diversity, indicating that the alteration of the structure of the bacterial community concerning its richness in several taxonomic groups is not significant for the studied cohort. However, there were significant differences in beta diversity, indicating a change in the diversity of some bacterial species between control dogs and those with YTE or in remission. There was no significant difference between YTE and Remission groups for beta diversity, although there are punctual significant differences in some species’ abundance. The dog microbiome is not that well characterised at the species level, indicating the need for additional characterisation of dog microbiome members that will facilitate future studies. Microbiome analysis of YTE dysbiosis showed a significant enrichment in bacterial ASVs belonging to taxa such as *Clostridium sensu stricto* 1, *Escherichia-Shigella, Streptococcus* and a significant depletion in ASVs belonging to taxa like *Bacteroides*, *Prevotella*, *Phascolarctobacterium*, *Fusobacterium* and *Alloprevotella*. Dogs that showed clinical improvements and were classified as in remission did not show a recovery from dysbiosis and were closer in bacterial composition to the YTE group than to the control one.

## Material and Methods

### Animals and sample collection

The study was approved by the Ethics Committee of the University of Veterinary Medicine Vienna and the Austrian Federal Ministry of Science and Research (BMWF-68.205/0150-V/3b/2018). All methods were carried out in accordance with relevant Austrian guidelines and regulations and the ARRIVE guidelines (https://arriveguidelines.org) were followed. Before entering the study, owners signed a written consent form. Client-owned Yorkshire Terriers (N=13) with chronic (≥3-week duration) or intermittent GI signs (diarrhea, vomiting, weight loss, anorexia) or pleura or abdominal effusion were presented to the Clinic for Small Animal Internal Medicine of the University of Veterinary Medicine Vienna, Austria, over a 30-month period (January 2019 and June 2021). These dogs were enrolled in the study as the YTE group. Diagnosis of YTE was made after exclusion of other possible causes of chronic GI symptoms and effusions and after histopathologic evidence of the characteristic intestinal inflammation pattern. Diagnostic tests included a physical exam, CBC, serum biochemical profile, measurement of bile acids and basal cortisol concentrations, ACTH-stimulation test (if basal cortisol < 2 μg/dl), and the assessment of serum concentration of cTLI (canine trypsin-like-immunoreactivity), SpecPL (specific pancreatic lipase), and cobalamin. Additionally, urinalysis for urine protein creatinine (UPC) ratio and urinary sediment, abdominal ultrasonography, and analysis of faecal samples by flotation and faecal Giardia antigen test were performed. All patients in the YTE group underwent gastroduodenoscopy at the time of presentation.

Accordingly, for the control group, adult client-owned healthy Yorkshire Terriers without any signs of GI disease (N=26) were prospectively enrolled. These dogs were defined as healthy after a diagnostic evaluation that included history, physical exam, CBC, chemistry profile, faecal analysis, urinalysis, and abdominal ultrasound. Dogs that had received treatment with antibiotics or glucocorticoid within the last 2 weeks and dogs that consumed a GI diet within the last two months were excluded.

Body conditioning scores (Nestle Purina: from 1 = very thin to 9 = significant obesity), faecal scores (Nestle Purina: from 1 = very firm to 7 = watery faeces), and muscle conditioning scores (WSAVA global nutrition committee: from A = normal muscle mass to D = severe muscle loss) were recorded. The canine chronic enteropathy activity index (CCECAI) was used to calculate clinical severity in all dogs. This scoring is based on severity alterations in 9 different IBD-relevant criteria. These criteria are: attitude and activity, appetite, vomiting, faecal consistency, defecation frequency, weight loss, serum albumin concentration, peripheral edema and ascites, and pruritus; each scored on a scale from 0-3. After summation, the total score is determined to be clinically insignificant (0–3), mild (4– 5), moderate (6–8), severe (9–11), or very severe (≥12) IBD^40^.

The score was calculated at enrolment day for dogs in both groups and additionally at the re-check for the dogs in the YTE group. A board-certified internist (in the author list: A.I.G.) performed all diagnostic investigations, gastroduodenoscopy and re-checks. One board-certified pathologist graded intestinal biopsies of the YTE dogs according to the World Small Animal Veterinary Association (WSAVA) and International Gastrointestinal Standardisation Group guidelines^41^. Dogs from the YTE group received a hydrolysed diet (Hill’s Prescription Diet z/d Canine) or a low-fat diet (Hill’s Prescription Diet i/d Low Fat Canine) in a double blindly design^42^. If the treatment results were inadequate after 14 days, diets were switched, and the dogs were revaluated after 28 days. The portion of food prescribed was aligned with the body weight recommendation of the manufacturer. After clinical remission was achieved (defined by a decrease of CCECAI scores to ≤3), the earliest 70 days after treatment initiation, individuals in the YTE group were revaluated, and faecal sampling was repeated. Due to ethical concerns, gastroduodenoscopy was not performed in control dogs, and not repeated in dogs in clinical remission.

### DNA extraction and 16S rRNA gene sequencing

DNA from stool samples was extracted with the Promega Maxwell® RSC Faecal Microbiome DNA Kit, and further prepared for amplicon sequencing in a two-step PCR as described previously^43^. The 515F (GTGYCAGCMGCCGCGGTAA)^44^ and 806R (GGACTACNVGGGTWTCTAAT)^45^ were used to amplify the V4 region of the 16S rRNA gene from most bacteria and archaea. Briefly, in the first amplification step, the V4 regions of the 16S rRNA genes were amplified with the 515F/806R primers. The PCR for each sample was set up in triplicate 25 μl reactions employing the DreamTaq Green PCR Master Mix (Thermo Fisher Scientific, USA), with 0.25 μmol L–1 of forward and reverse primer each and 2 μl DNA template. Four PCR negative controls (PCR with nuclease-free water as template) were routinely performed. The PCR products were subsequently analysed by gel electrophoresis. Thereafter, triplicate first-step PCR reactions were pooled and purified and normalised with the SequalPrep Normalisation Plate Kit (Invitrogen) following the manufacturers’ instructions. The PCR products were combined with a kit-specific binding buffer and transferred into a SequelPrep normalisation plate, the walls of which are coated with a DNA-binding solid phase capable of retaining a maximum of 25 ng DNA. Unbound PCR products (including short fragments, which cannot bind to the solid phase) and buffer were removed, and the normalised PCR products were detached from the DNA-binding solid phase with a kit-specific elution buffer.

The second amplification step (barcoding reaction) consisted of two barcoding primers. The forward primer had 12 nucleotide barcode while the reverse one used 16 nucleotide head (*5*’-BC12_1-H1-*3’* and *5*’-BC12_2-H2-*3*’) for amplification. The purchased oligonucleotide barcodes used for this study were synthesised and HPLC purified by Microsynth (Balgach, Switzerland). Barcoding PCR reactions were set up as a single 50 μl reaction using the DreamTaq PCR Master Mix (Thermo Fisher Scientific, USA), with 0.8 μmoles of each used barcoding primer and 10 μL of purified and normalised first-step PCR product as template. The barcoding PCR was performed with following thermal cycle conditions: initial denaturation at 94°C for 4 min; 7 cycles of denaturation at 94°C for 30 s, annealing at 52°C for 30 s, and elongation 72°C for 60 s; followed by a final elongation step at 72°C for 7 min. Thereafter, samples were again purified and normalised with the SequalPrep Normalisation Plate Kit as described above.

The amplicon pool fragment size was verified on a D1000 ScreenTape Assay using a TapeStation 4200 (Agilent), and the DNA concentration in the amplicon pool was quantified with a Quant-iT dsDNA Assay (ThermoFisher) on a Qubit 4 Fluorometer (ThermoFisher). Sequencing libraries were prepared and indexed by adapter ligation and PCR (8 cycles) using the TruSeq Nano DNA Library Prep Kit (Illumina) and TruSeq DNA Single Indexes Set A or B (Illumina), excluding the DNA fragmentation and clean-up of fragmented DNA steps, and otherwise following the manufacturer’s instructions. Sequencing library size and the efficiency of unligated adaptor removal were verified on a D1000 ScreenTape Assay using a TapeStation 4200, and the sequencing library concentration was determined with a Quant-iT dsDNA Assay on a Qubit 4 Fluorometer. For each sequencing run, 2–5 amplicon pool libraries (in total up to 1,092 distinct amplicon libraries) were combined and sequenced in 2 × 300 cycle paired-end mode on an Illumina MiSeq using the MiSeq Reagent kit v3 (Illumina) with 6 nucleotides library indexes (DNA Single Indexes Set A or B, Illumina). To achieve sufficient variability during the first five sequencing cycles, which is necessary for efficient sequence cluster identification and phasing/prephasing calibration during Illumina sequencing, we spiked in 1–7 random shotgun genomic or metagenomic sequencing libraries (to an abundance of 9–21%) and 1% PhiX control to each sequencing run. As denoising methods, we opted for the use of the <u>A</u>mplicon <u>S</u>equence <u>V</u>ariants (ASV) since it has a better performance than other methods such as the use of <u>O</u>perational <u>T</u>axonomic <u>U</u>nits (OTU) when it comes to estimate microbial biodiversity^46^. Amplicon sequence variants (ASVs) were inferred using the DADA2 R package^47^ applying the recommended workflow^48^. FASTQ reads 1 and 2 were trimmed at 220 nucleotide and 150 nucleotide with allowed expected errors of 2, respectively. ASV sequences were subsequently classified using DADA2 and the SILVA database SSU Ref NR 99 release 138.1^49,50^ using a confidence threshold of 0.5. ASVs classified as mitochondria or chloroplast were removed. Only samples sequenced with at least 5000 read pairs were retained for downstream analyses.

### Statistical analysis

Differences in the canine chronic enteropathy activity index (CCECAI) among groups were inferred from a Kruskal-Wallis test followed by Mann-Whitney U-test to compare the three groups among them. Alpha diversity metrics, Bray–Curtis dissimilarity, and the principal analysis components were calculated using the R package ampvis2 version 2.6.5^51^ with the R package vegan version 2.5–6^52^. Alpha and beta diversity indexes were performed on species level. Alpha diversity data were rarified to 6793 counts. In the case of beta diversity we used non-rarified data with Aitchison distance, implying that the normalisation method was centered log-ratios. All statistical comparisons between groups were performed using an ANOVA and in case of significant difference, the ANOVA was followed by a post-hoc test Tukey HSD in R. Adjusted *P*-value significance codes used in all figures are *<0.05, **<0.01, ***<0.001, and ****<0.0001. The difference in per-group centroids was tested with a PERMANOVA on Aitchison distance using vegan and microViz^53^. All PERMANOVAs were strictly on one category, with no controls for potential co-variates.

## Data Availability

The datasets generated and/or analysed during the current study are available in the NCBI BioProject repository with the accession number PRJNA861113 and can be directly accessed with the following link: https://www.ncbi.nlm.nih.gov/bioproject/PRJNA861113/.

## Funding

This research was partially funded by Hill’s Clinical Study Grant. The authors acknowledge financial support from European Union’s Horizon 2020 research and innovation program under the Marie Sklodowska-Curie grant agreement No. 898858 and European Union’s Horizon 2020 research and innovation program under grant agreement No. 857287. Also the Joint Microbiome Facility of the Medical University of Vienna and the University of Vienna (project ID JMF-2101-12) contributed to this work.

## Acknowledgements

We would like to thank Dr Petra Pjevac from the Joint Microbiome Facility of the Medical University of Vienna and the University of Vienna for useful discussions and assistance. We are also grateful to Dr Barbara Pratscher (from Small Animal Internal Medicine, University of Veterinary Medicine, Vienna) for comments and suggestions.

## Notes

### Competing Interest Statement

The authors have declared no competing interest.

### Summary of Updates

The text has been proofread and the discussion has been extended.

