## Supplementary tables and figures for "Gut Microbiome Signatures of Yorkshire Terrier Enteropathy during Disease and Remission": Fig S1.pdf

**Supplementary Material**

**Fig. S1. Principal Component Analysis of clinical and clinicopathological parameters does not segregate the three studied groups (YTE, Remission and Control).** The test included as parameters were complete blood count (CBC), serum biochemical profile, measurement of bile acids and basal cortisol concentrations, serum concentration of cTLI (canine trypsin-like- immunereactivity), SpecPL (specific pancreatic lipase), and cobalamin. Additionally, urinalysis for urine protein creatinine ratio (UPC) and urinary sediments.

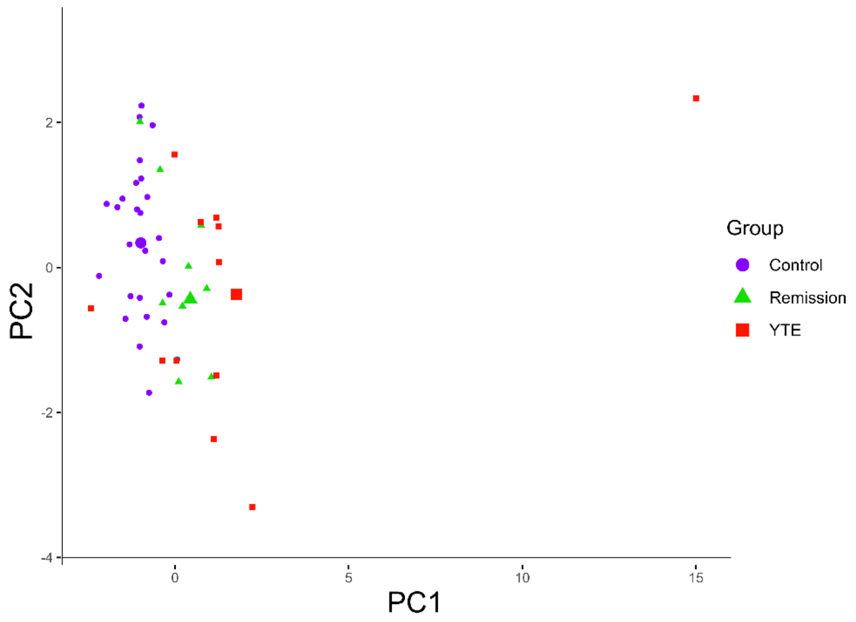
