## Supplementary tables and figures for "Gut Microbiome Signatures of Yorkshire Terrier Enteropathy during Disease and Remission": Fig S2 .pdf

**Fig. S2.** Analysis of the differentially abundant fraction expressed as numbers of reads of 16S rRNA gene for individual dogs and the taxonomic breakdown of the dominant ASVs across all samples.

JMF-2101-12 | gene: 16S rRNA V4 | state: refined | filter min count: 5000 | agglomerated by rank: Domain

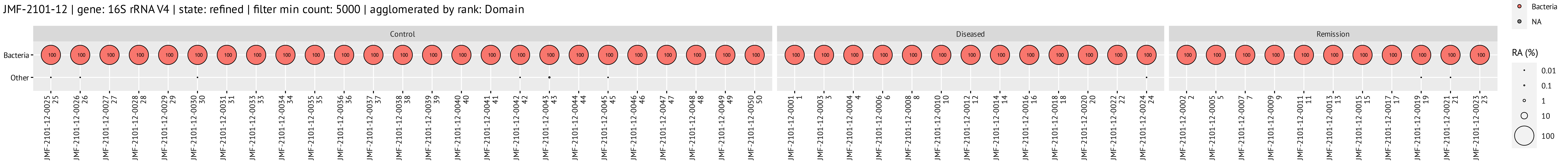


Prevalence collapse settings: "Others" except ≥1% RA in at least 3 samples.

JMF-2101-12 | gene: 16S rRNA V4 | state: refined | filter min count: 5000 | agglomerated by rank: Phylum

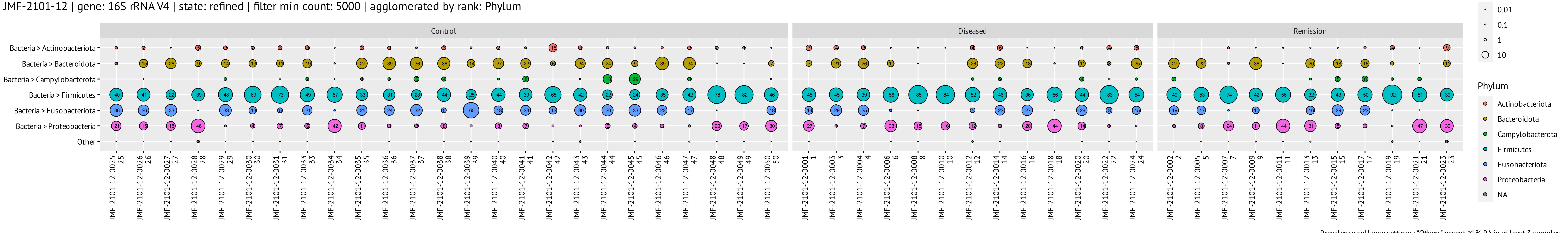


Prevalence collapse settings: "Others" except ≥1% RA in at least 3 samples.

JMF-2101-12 | gene: 16S rRNA V4 | state: refined | filter min count: 5000 | agglomerated by rank: Class

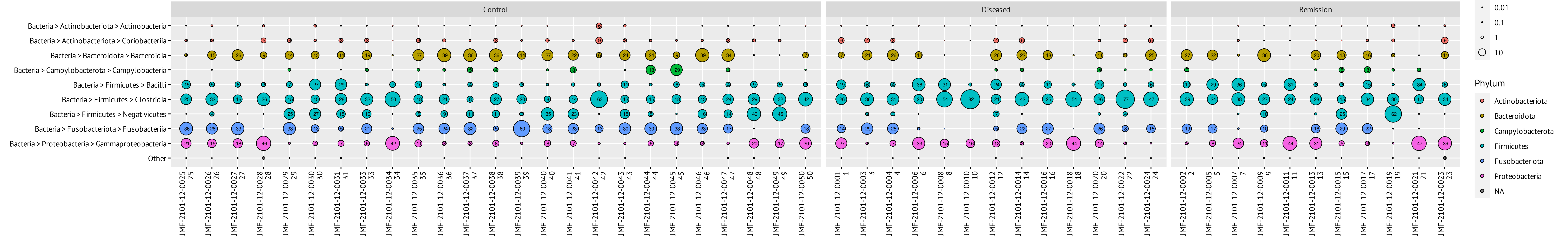


Prevalence collapse settings: "Others" except ≥1% RA in at least 3 samples.

JMF-2101-12 | gene: 16S rRNA V4 | state: refined | filter min count: 5000 | agglomerated by rank: Order

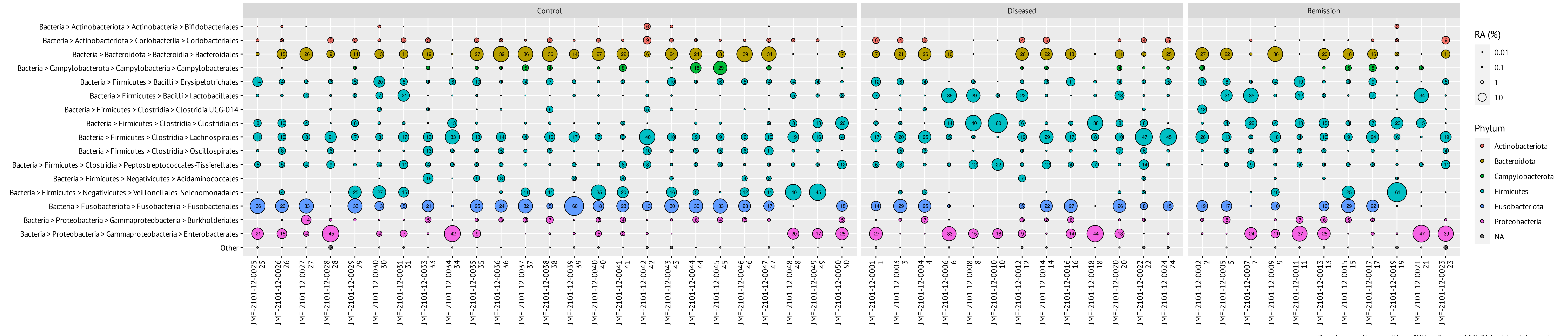


Prevalence collapse settings: "Others" except ≥1% RA in at least 3 samples.

JMF-2101-12 | gene: 16S rRNA V4 | state: refined | filter min count: 5000 | agglomerated by rank: Family

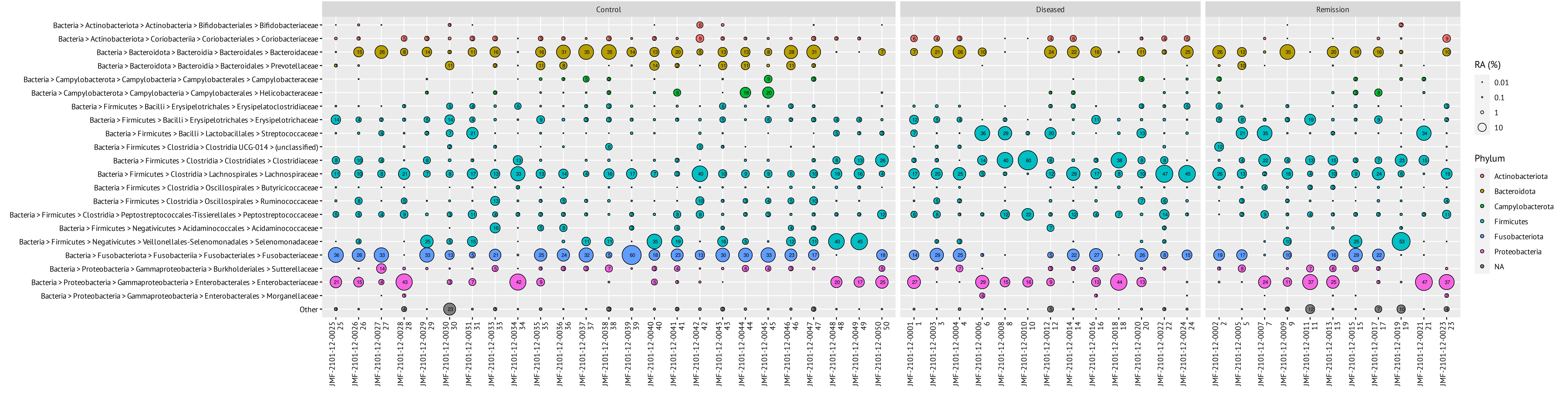


Prevalence collapse settings: "Others" except ≥1% RA in at least 3 samples.

JMF-2101-12 | gene: 16S rRNA V4 | state: refined | filter min count: 5000 | agglomerated by rank: Genus

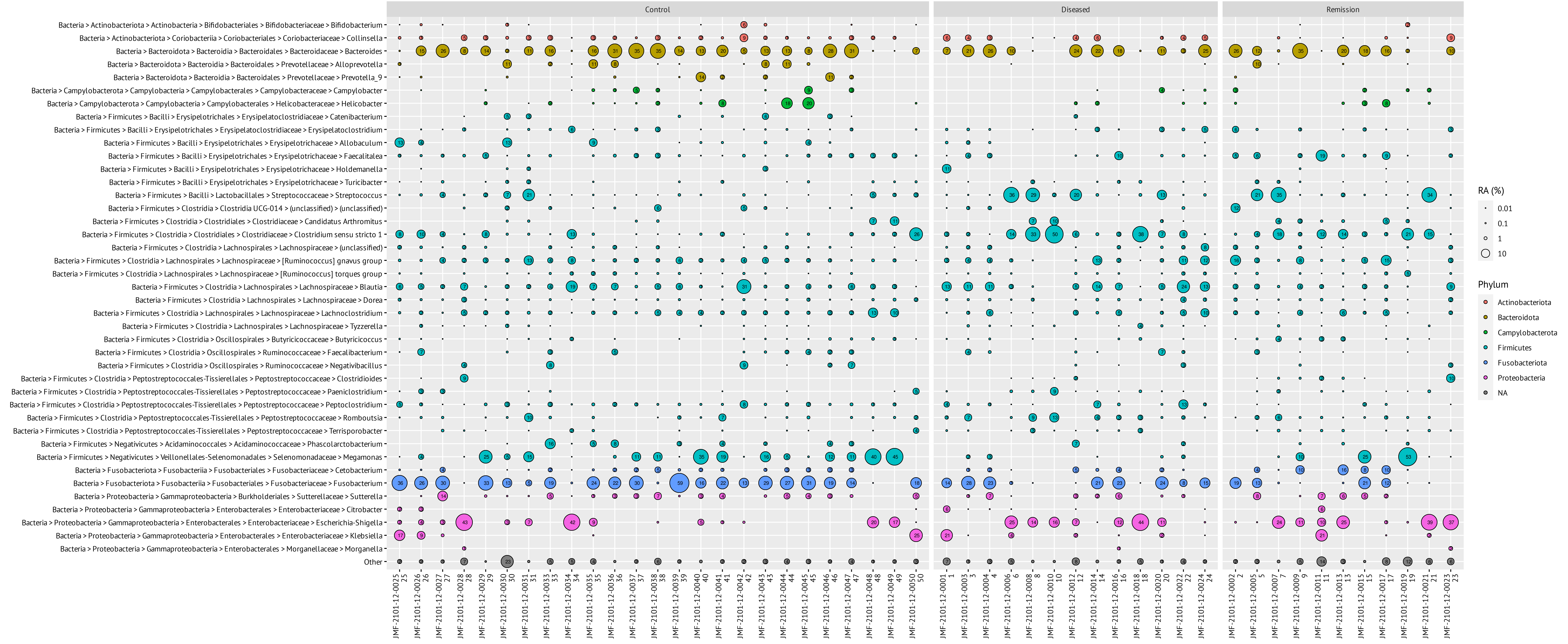


Prevalence collapse settings: "Others" except ≥1% RA in at least 3 samples.

JMF-2101-12 | gene: 16S rRNA V4 | state: refined | filter min count: 5000 | agglomerated by rank: Species

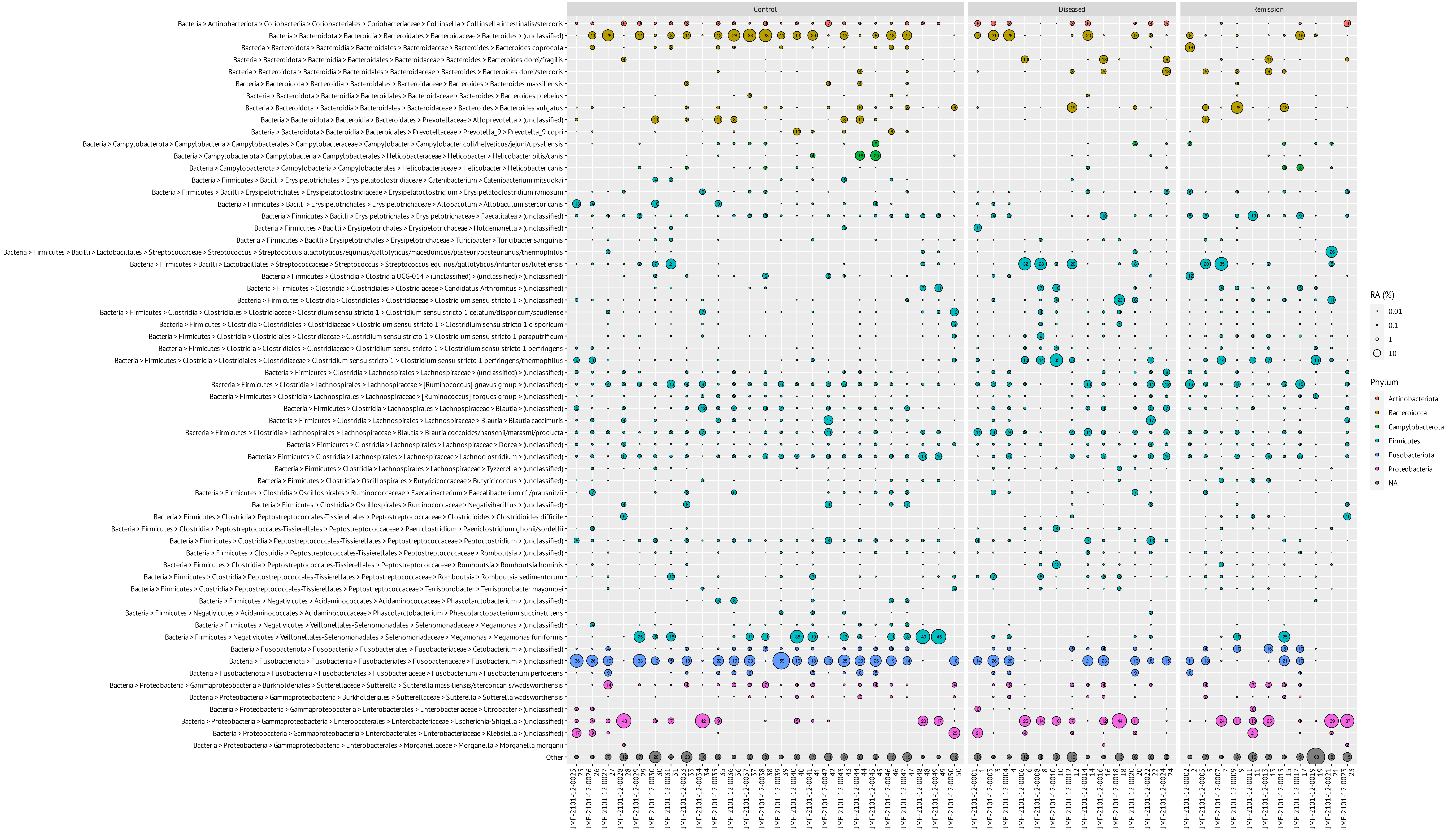


Prevalence collapse settings: "Others" except ≥1% RA in at least 3 samples.

JMF-2101-12 | gene: 16S rRNA V4 | state: refined | filter min count: 5000

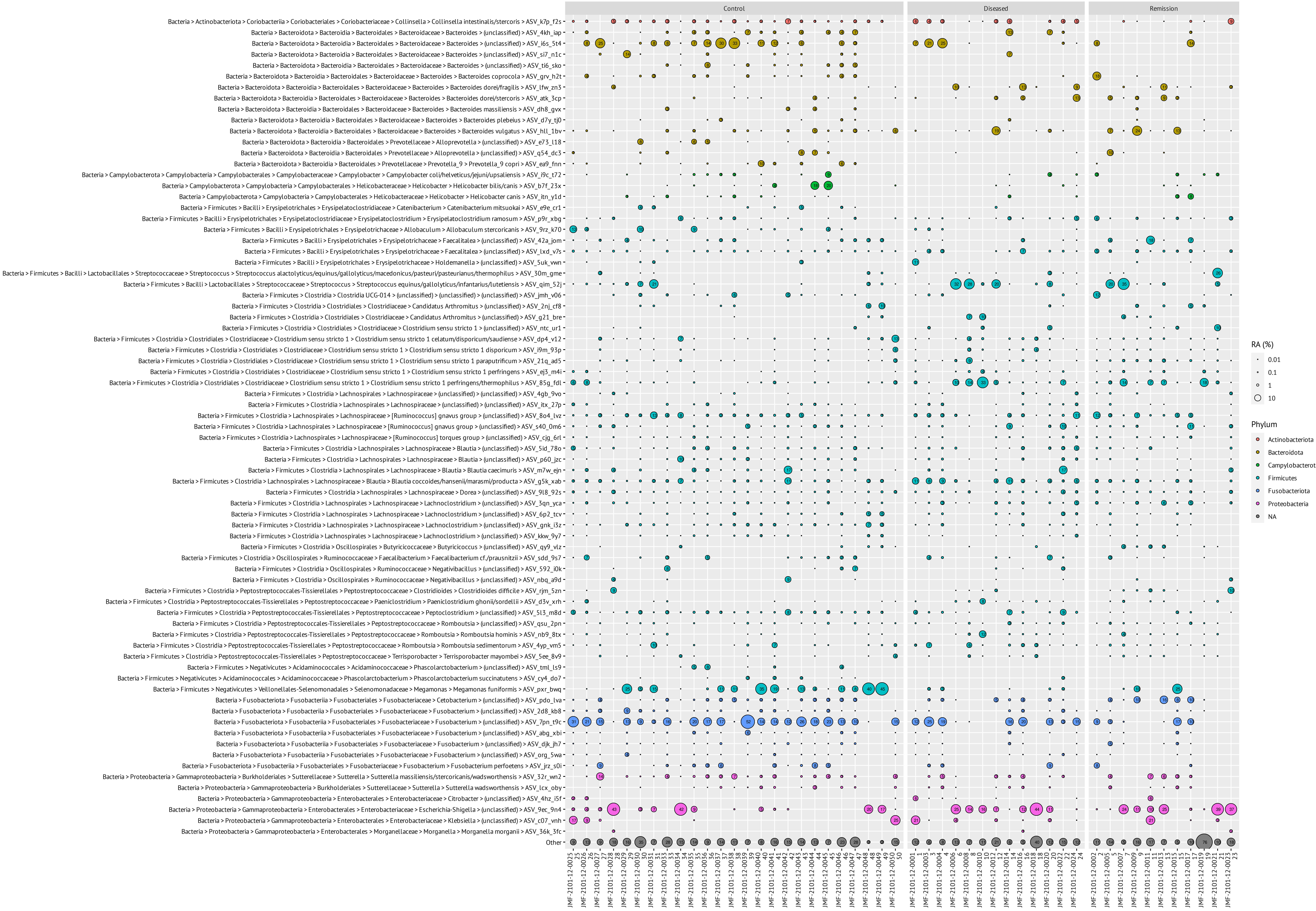


Prevalence collapse settings: "Others" except ≥1% RA in at least 3 samples.
