## Supplementary figures and images for "Gut Microbiome Signatures of Yorkshire Terrier Enteropathy during Disease and Remission"

### Fig 1.tif

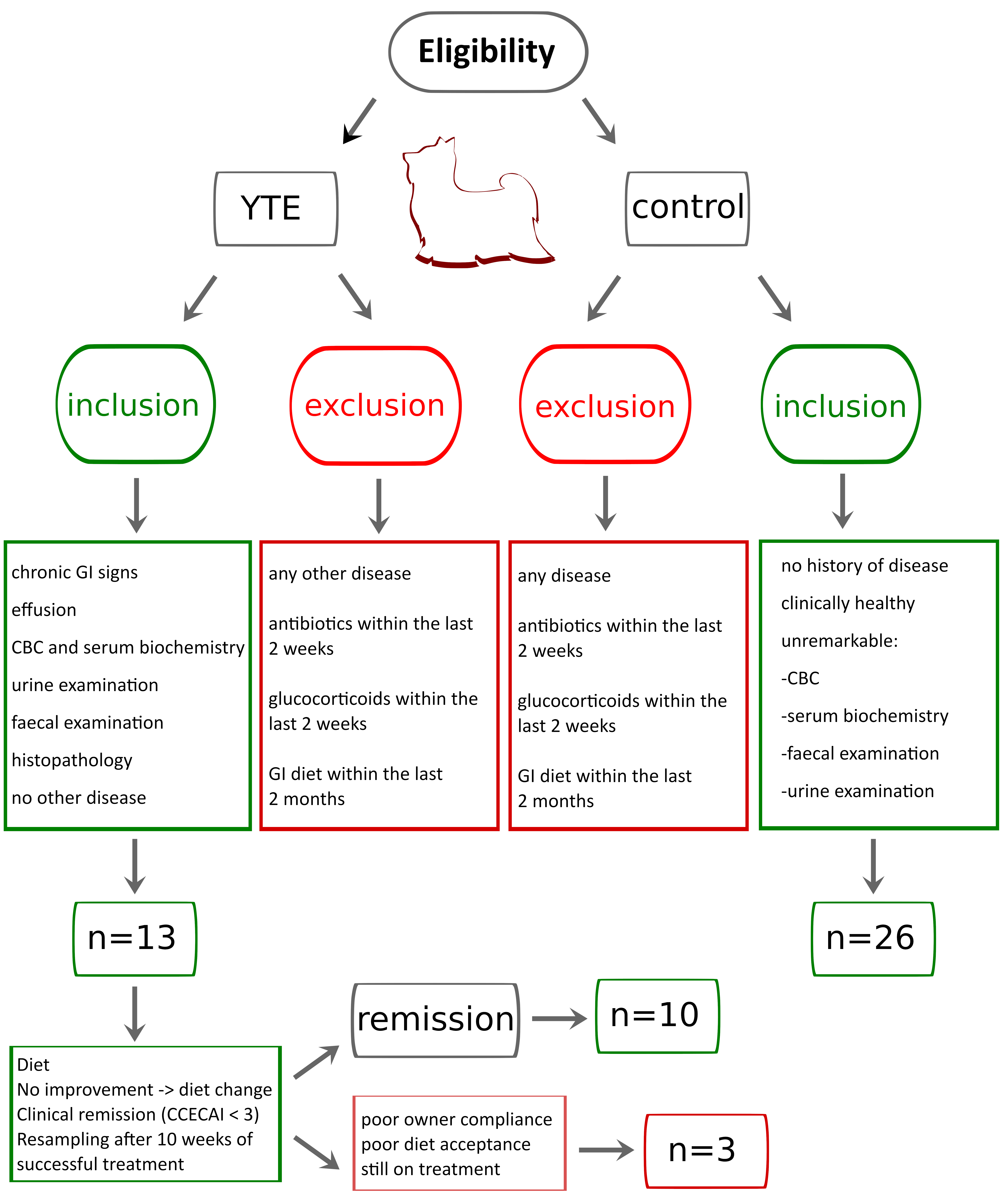

### Figure 2.tif

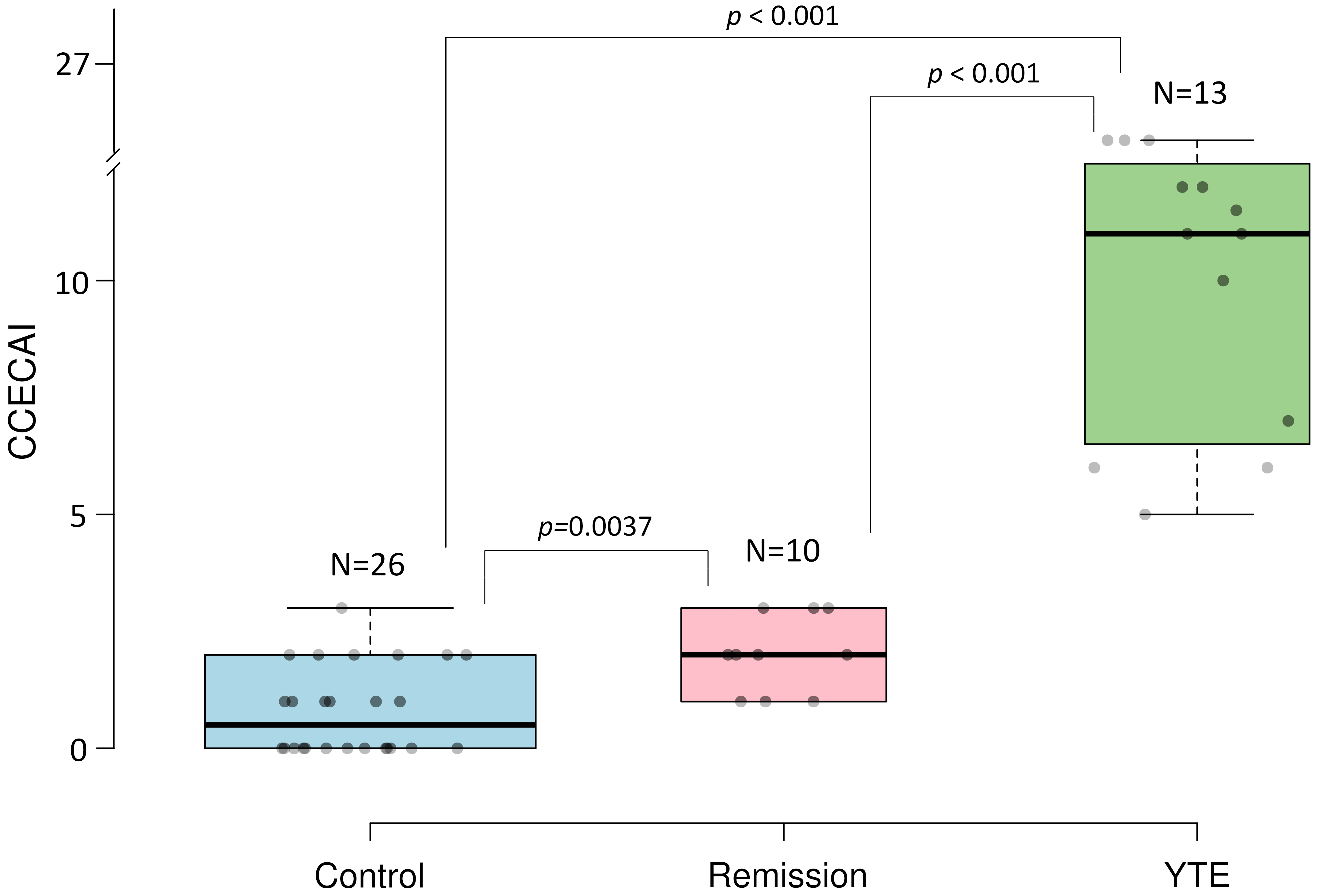

### Figure 3.tif

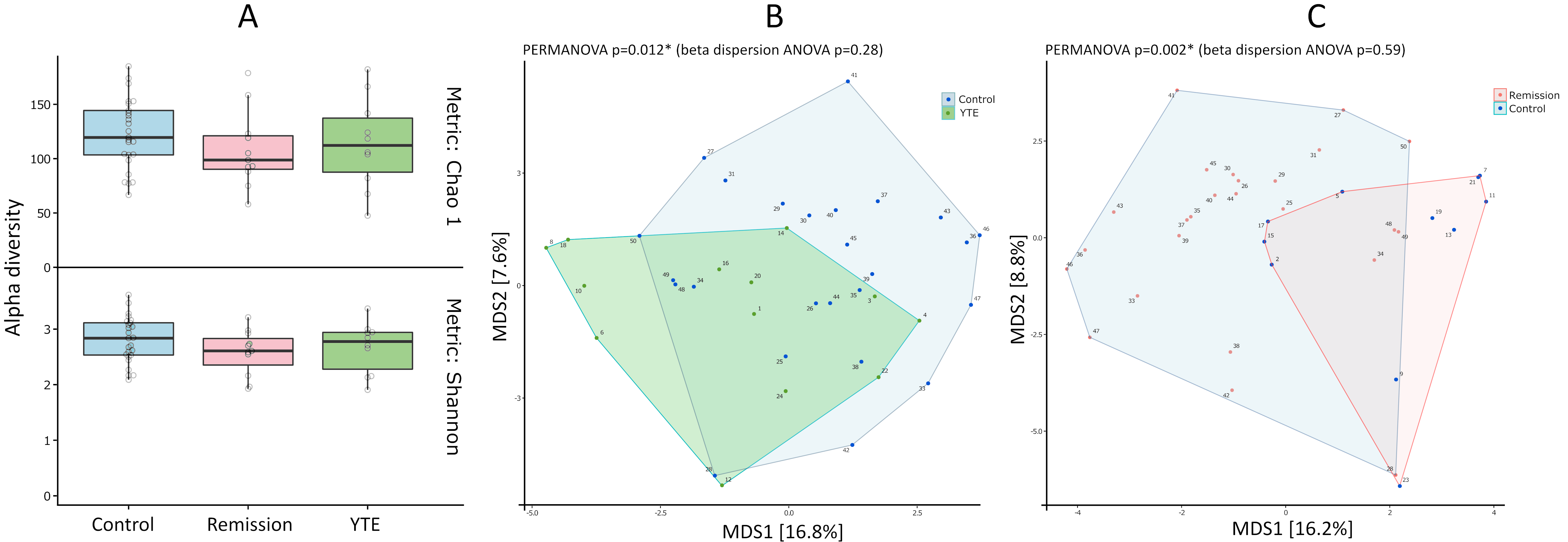

### Figure 4.tif

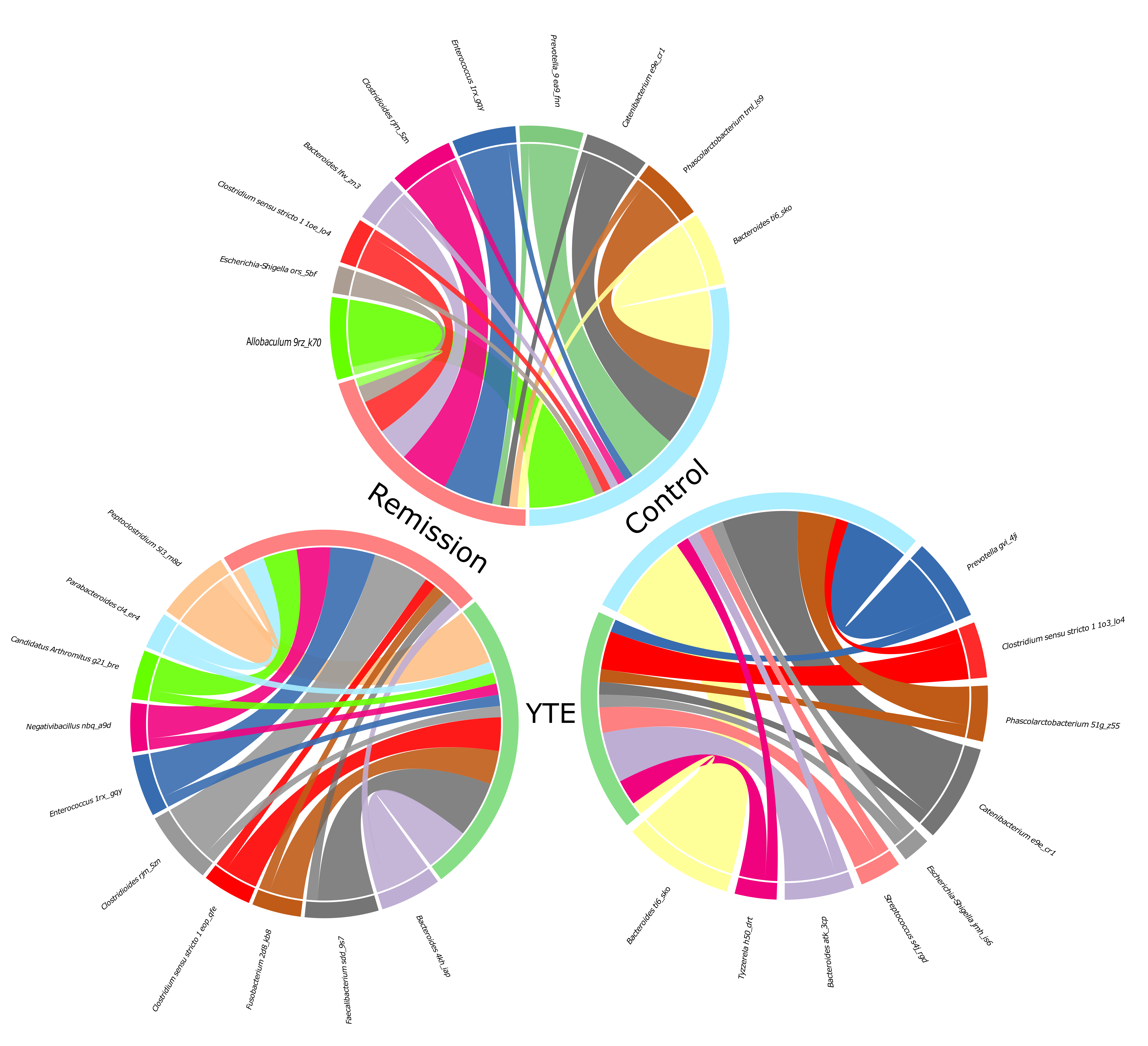

### Figure 5.tif

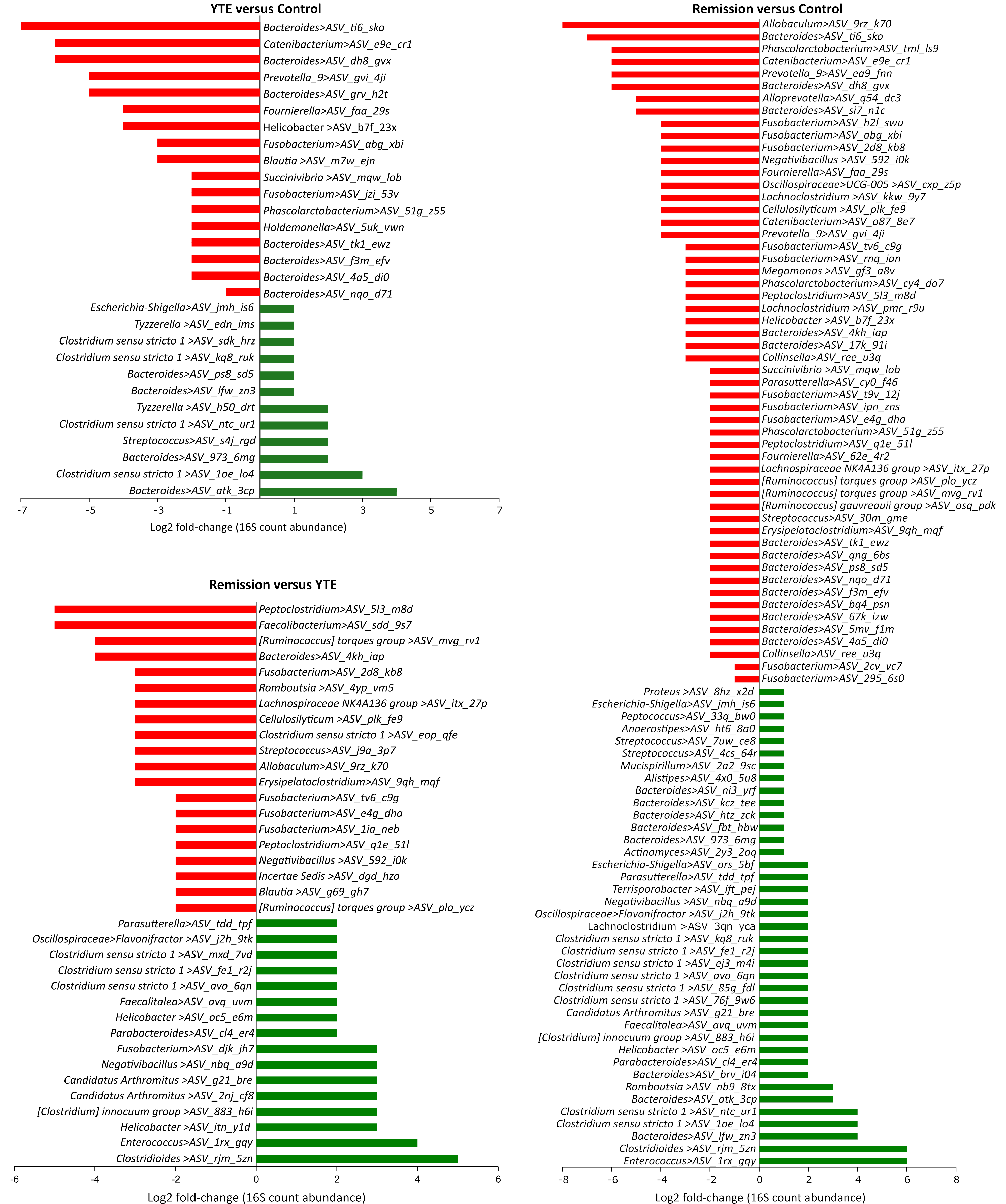
